# Elevational patterns in plant mating systems and pollen limitation in Afrotropical montane grasslands

**DOI:** 10.64898/2026.01.09.698470

**Authors:** Dominik Anýž, Štěpán Janeček, Shedrach B. Kongvong, Carlson T. Bawe, Petr Chlup, Sylvain Delabye, Nestoral T. Fominka, Fernando P. Gaona, Jiří Hodeček, Petra Janečková, Ishmeal N. Kobe, Francis E. Luma, Ondřej Mottl, Sailee P. Sakhalkar, Robert Tropek

**Affiliations:** Department of Ecology, Faculty of Science, Charles University, Prague, Czechia; A.P. Leventis Ornithological Research Institute, University of Jos, Jos, Nigeria; Faculty of Science, University of Buea, Buea, Cameroon; Department of Forestry and Wildlife, Faculty of Agriculture and Veterinary Medicine, University of Buea, Buea, Cameroon; Institute of Entomology, Biology Centre, Czech Academy of Science, České Budějovice, Czechia; Swiss Human Institute of Forensic Taphonomy, University Centre of Legal Medicine Lausanne-Geneva, Lausanne; Department of Botany, Faculty of Science, University of South Bohemia, Czechia; Bokwaongo Village Forest Management Committee (BVFMC), Bokwaongo, Cameroon; Mount Cameroon National Park, Buea, Cameroon; Department of Botany, Faculty of Science, Charles University, Prague, Czechia; Center for Theoretical Study, Charles University, Prague, Czechia; Laboratory of Environmental Microbiology, Institute of Microbiology of the Czech Academy of Sciences, Prague, Czechia

**Keywords:** altitudinal gradients, pollination biology, plant breeding systems, self-pollination, self-compatibility, tropical mountains

## Abstract

1. Plant mating systems and pollen limitation often vary along elevational gradients, yet empirical evidence remains mixed and rarely comes from multi-species experimental studies in tropical mountain ecosystems.
2. We tested how elevation affects natural seed set, pollen limitation, and selfing capacity, and whether these patterns are associated with variation in flower visitation, in Afromontane grasslands on Mount Cameroon. We conducted a hand-pollination experiment on seven zoogamous plant species at four elevations above the timberline (2,300–3,800 m), applying autonomous selfing, geitonogamous selfing, outcrossing, and open-pollination treatments to 1,776 flowers and quantifying seed set. In parallel, we quantified pollinator visitation on unmanipulated plants to estimate visitation frequency, morphospecies richness, and functional-group richness.
3. Natural reproductive success of the studied plants exhibited a pronounced mid-elevation peak and declined sharply towards the summit, where pollen limitation increased strongly. At the highest elevation, some species produced few or no seeds even under outcross-pollination, indicating physiological reproductive constraints.
4. Indices of autonomous selfing and geitonogamy varied among species, with no consistent elevational patterns. Despite detected partial self-compatibility, summit populations did not exhibit expected shifts towards selfing, suggesting limited reproductive assurance under high-elevation conditions.
5. Pollinator visitation frequency and diversity declined at the highest elevation. Natural seed set was positively associated with visitation frequency and mildly negatively associated with morphospecies richness, whereas pollen limitation and selfing indices showed no clear relationships with visitation metrics, consistent with the influence of additional physiological and developmental constraints on reproduction.
6. **Synthesis.** Our study shows that in isolated Afrotropical montane grasslands, plant reproductive success at high elevations is jointly constrained by declining pollination service and abiotic limitations, which is not compensated by increased selfing. This suggests that successful plant reproduction near upper vegetation limits depends on both sufficient pollination service and favourable physiological conditions. Under ongoing climate change, upslope range shifts of plants may therefore not guarantee reproductive success if plant and pollinator responses to warming are asynchronous or if a part of extreme high-elevation conditions remain limiting. These findings advance understanding of how biotic interactions and abiotic constraints together shape plant reproduction along tropical elevational gradients.

## Introduction

The balance between outcrossing and selfing is a central determinant of reproductive success across ecological gradients in flowering plants. Although some plants are obligately outcrossing and others predominantly selfing, most angiosperms function mainly as outcrossers and depend on compatible cross-pollen to maintain genetic diversity (Ferrer and Good-Avila, 2007; Wright et al., 2013; Ferrer et al., 2025). At the same time, numerous species exhibit partial or full self-compatibility, enabling autonomous selfing within flowers (autogamy) or facilitated selfing among flowers of the same individual (geitonogamy), particularly when pollinator service is unreliable, constrained, or seasonally variable (Dafni et al., 1995; Takebayashi and Morrell, 2001; Willmer, 2011). The interplay between these strategies is strongly influenced by abiotic stress and pollinator availability, linking plant reproductive systems directly to environmental conditions (Knight et al., 2005; García-Camacho and Totland, 2009).

Elevational gradients impose sharp changes in temperature, wind exposure, solar radiation, and growing-season duration over short distances, with expectable effects on flowering phenology, pollen transfer, and seed production (Arroyo et al., 1982; McCall and Primack, 1992; Blionis et al., 2001). Harsh climatic conditions at high elevations typically reduce pollinator abundance, taxonomic and functional diversity, and the duration of daily foraging activity, thereby constraining animal-mediated pollen transfer (Arroyo et al., 1985; McCall and Primack, 1992; Totland, 1997; Medan et al., 2002). These reductions in pollinator service are frequently associated with increased pollen limitation, i.e. reduced reproductive success due to insufficient pollination, in montane communities (Knight et al., 2005; García-Camacho and Totland, 2009; Wu et al., 2019; Jiang and Xie, 2020). Plants may respond through multiple, often concurrent, strategies, including extending floral longevity to increase the probability of pollination during short favourable periods (Bingham and Orthner, 1998; Blionis et al., 2001) or shifting mating systems towards a higher contribution of selfing (Herlihy and Eckert, 2002; Kalisz and Vogler, 2003). In this context, both interspecific and intraspecific variation in partial self-compatibility and selfing rates represent key evolutionary strategies by which plant populations respond to spatiotemporal limitations in pollinator service (Schoen et al., 1996; Busch and Delph, 2012; Barrett et al., 2014). Nevertheless, our understanding of how plants adjust their reproductive strategies along elevational gradients remains limited by the scarcity of experimental studies quantifying intraspecific variation across broader elevation ranges, particularly in tropical ecosystems (Wu et al., 2019; Jiang and Xie, 2020).

Empirical studies along elevational gradients provide mixed support for the expectation that selfing increases with rising elevations. Interspecific comparisons often report a higher prevalence of selfing or a shift towards wind pollination among high-elevation floras (Medan et al., 2002; Berry and Calvo, 2016), consistent with strong pollen limitation in alpine habitats (Jiang and Xie, 2020). Intraspecific patterns, however, are considerably more variable: some species show increased selfing rates in alpine populations (Etcheverry et al., 2008; Seguí et al., 2018; Abdusalam et al., 2020), others show little or no change (Gugerli, 1998; Young et al., 2002; Dai et al., 2017; Black et al., 2019), and some even exhibit reduced selfing at higher elevations (Wirth et al., 2010). Such recurrent findings of the increased selfing with elevation tend to occur in species already capable of autonomous self-pollination at lower sites (Arroyo et al., 2006; Pérez et al., 2013), suggesting that elevational trends frequently represent quantitative shifts in partial self-compatibility rather than transitions between outcrossing and selfing. However, much of this work is either taxonomically narrow or based on single-elevation snapshots of breeding systems, and relatively few studies test intraspecific changes in selfing capacity and pollen limitation across multiple elevations using comparable experimental protocols. In addition, studies that simultaneously manipulate pollen receipt and quantify pollinator visitation are still uncommon, particularly in tropical mountains and in Africa.

To investigate how elevational changes in environmental conditions and pollinator communities shape plant reproductive strategies in tropical Afromontane grasslands, we conducted a standardized pollination experiment on seven zoogamous species above the timberline on Mount Cameroon. Combining hand pollination with video recording of floral visitors, we quantified natural seed set, pollen limitation, and indices of autonomous and facilitated selfing (autogamy and geitonogamy), together with visitation frequency and the taxonomic and functional richness of floral visitors. Specifically, we asked (i) whether natural seed set, pollen limitation, and selfing show consistent intraspecific patterns along elevation and (ii) whether these elevational patterns can be explained by corresponding variation in pollinator visitation and diversity. We expected reduced natural seed set, increased pollen limitation, and stronger reliance on selfing at higher elevations, driven by the expected decline in pollinator activity and diversity under colder and generally harsher high-elevation conditions.

## Methods

### Study area and sites

This study was conducted on Mount Cameroon (4.203°N, 9.170°E), an active volcano rising to 4,040 m a.s.l. and forming the highest peak in West and Central Africa. The mountain represents a major biodiversity hotspot, harbouring numerous endemic species of plants and animals (Bergl et al., 2007; Ustjuzhanin et al., 2018; Janeček and Janečková, 2025). Its steep elevational gradient supports a wide range of habitats from lowland rainforest to subalpine heathlands. We focused on the Afromontane grasslands located above the timberline, which averages at ∼2,200 m a.s.l., i.e. lower than is typical for this region due to volcanic and anthropogenic disturbance (Cable and Cheek, 1998; Doležal et al., 2022).

We established four study sites at approximately 2,300, 2,800, 3,400, and 3,800 m a.s.l., spanning the full elevational range of Afromontane grasslands on Mount Cameroon (Fig. 1). The 2,300 m site was located along the Bakingili trail, and the three higher sites along the Guinness trail, all selected to maximise habitat representativeness, accessibility, and the diversity and abundance of flowering plants, while avoiding obviously degraded areas. Together, they cover the transition from species-rich montane grasslands with scattered shrubs to sparse herbaceous vegetation of the summit zone (Janeček and Janečková, 2025).

**Figure 1.**
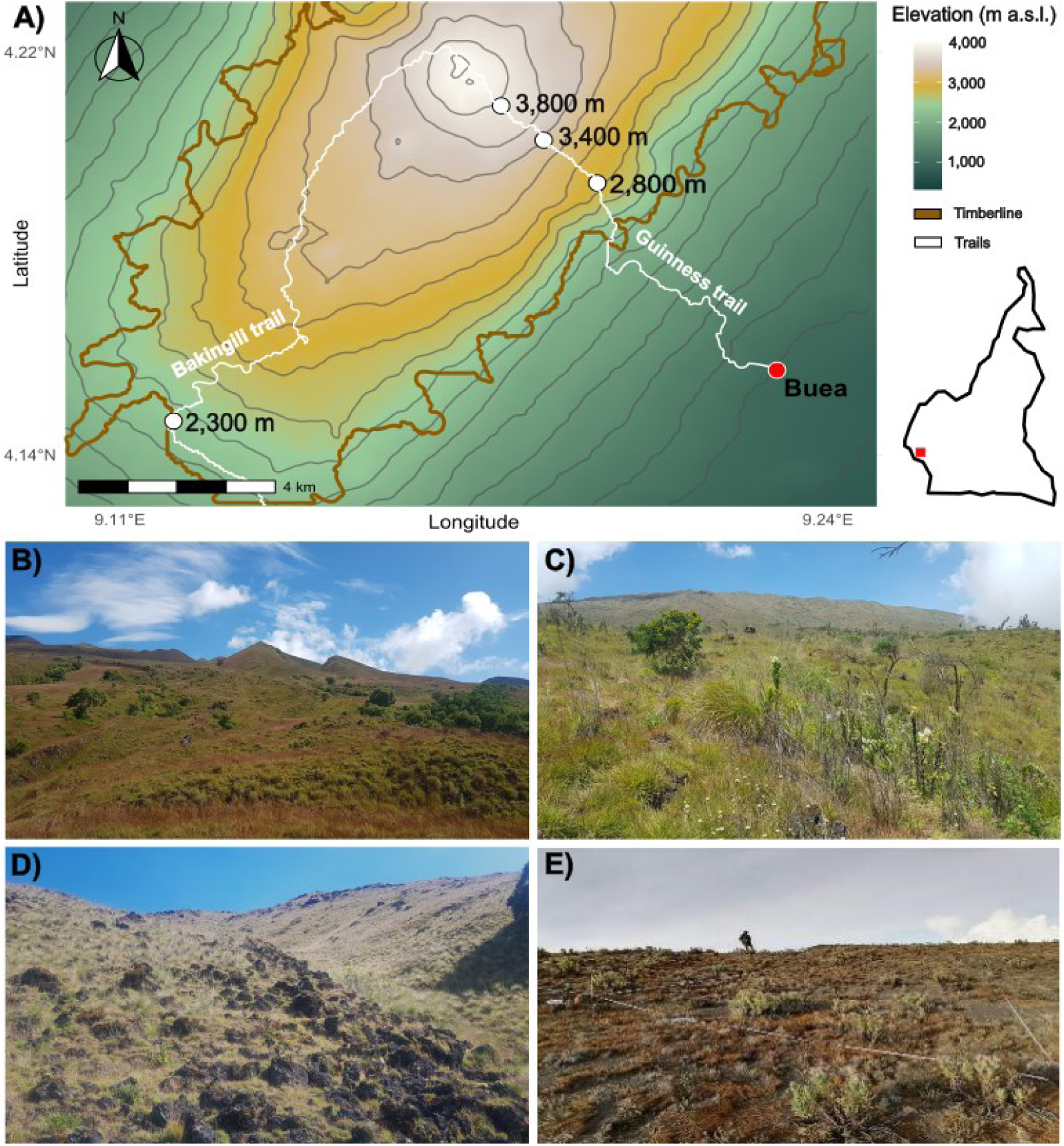
Study sites in Afromontane grasslands along the elevational gradient on Mount Cameroon. A) Shaded relief map of the upper south and southwestern slopes showing the four study sites at 2,300, 2,800, 3,400, and 3,800 m a.s.l. (white circles). Background colours indicate elevation, the brown line marks the timberline, and white lines show the main access trails. The inset map shows the position of Mount Cameroon (red dot) within Cameroon. B–E) Habitat structure at the four studied elevations: B) 2,300 m, C) 2,800 m, D) 3,400 m, and E) 3,800 m a.s.l., from relatively tall, shrub-rich grasslands at lower elevations to sparse, low herb cover near the summit. Credits: Dominik Anýž (A-D), Ishmeal N. Kobe (E).

### Pollination experiment

Seven plant species with expected zoogamy (Table 1) were selected for the experiment, based on their sufficient abundance across at least two elevations and floral morphology suitable for hand pollination. Within each site, several experimental patches were chosen where plants were accessible, abundant, and protected from fire. Within these patches experimental plant specimens were selected based on the availability of flower buds; the number of individuals and flowers treated varied among species and elevations according to their local abundance and flowering synchrony (Table 1).

**Table 1.**
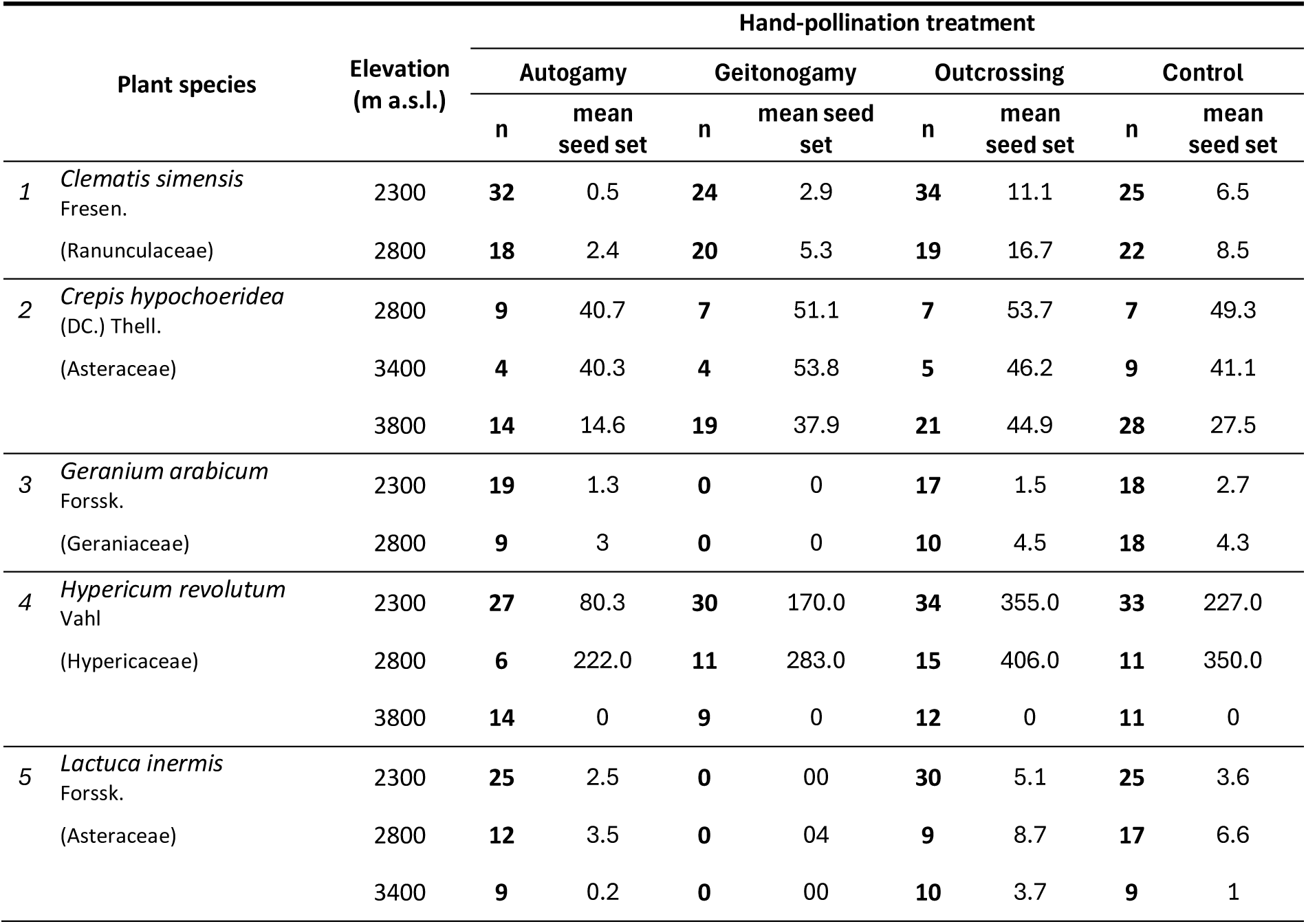
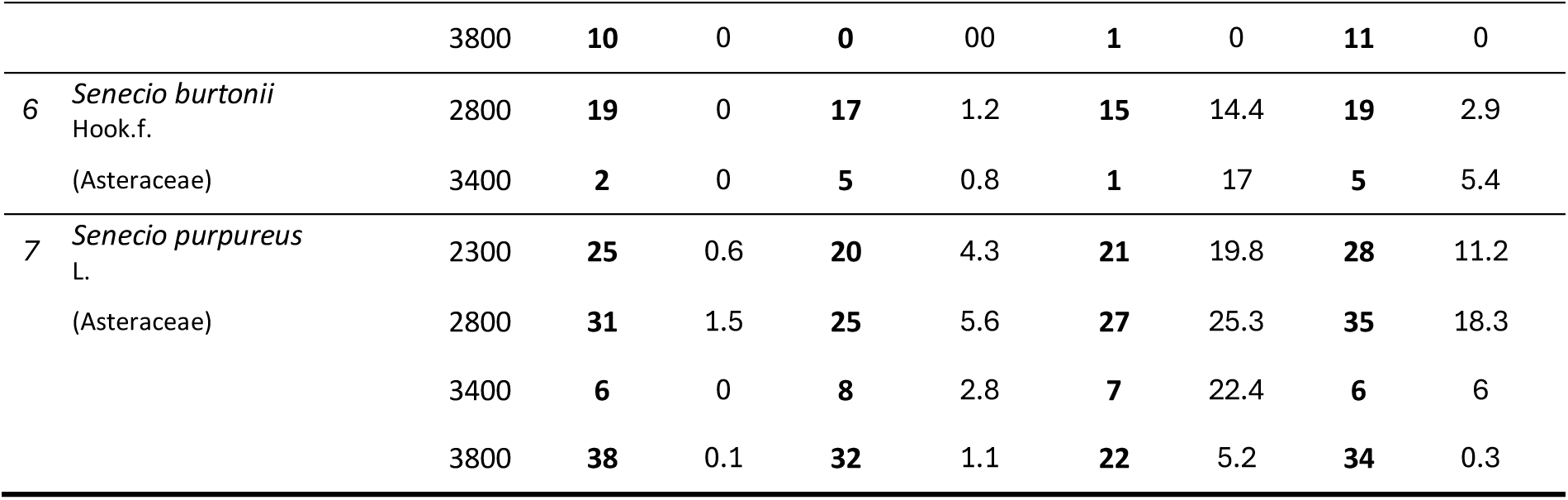
Plant species included in the pollination experiment, with the number of treated flowers (n) and mean seed set produced under each hand-pollination treatment at each elevation on Mount Cameroon. Seed set represents the number of mature seeds per treated flower (or per capitulum in composite species). Nomenclature follows Janeček and Janečková (2025).

| Plant species | Elevation<br>(m a.s.l.) | Hand-pollination treatment |  |  |  |  |  |  |  |
| --- | --- | --- | --- | --- | --- | --- | --- | --- | --- |
|  |  | Autogamy |  | Geitonogamy |  | Outcrossing |  | Control |  |
|  |  | n | mean<br>seed set | n | mean seed<br>set | n | mean<br>seed set | n | mean<br>seed set |
| 1 *Clematis simensis*<br>Fresen.<br>(Ranunculaceae) | 2300 | 32 | 0.5 | 24 | 2.9 | 34 | 11.1 | 25 | 6.5 |
|  | 2800 | 18 | 2.4 | 20 | 5.3 | 19 | 16.7 | 22 | 8.5 |
| 2 *Crepis hypchoeridea*<br>(DC.) Thell.<br>(Asteraceae) | 2800 | 9 | 40.7 | 7 | 51.1 | 7 | 53.7 | 7 | 49.3 |
|  | 3400 | 4 | 40.3 | 4 | 53.8 | 5 | 46.2 | 9 | 41.1 |
|  | 3800 | 14 | 14.6 | 19 | 37.9 | 21 | 44.9 | 28 | 27.5 |
| 3 *Geranium arabicum*<br>Forssk.<br>(Geraniaceae) | 2300 | 19 | 1.3 | 0 | 0 | 17 | 1.5 | 18 | 2.7 |
|  | 2800 | 9 | 3 | 0 | 0 | 10 | 4.5 | 18 | 4.3 |
| 4 *Hypericum revolutum*<br>Vahl<br>(Hypericaceae) | 2300 | 27 | 80.3 | 30 | 170.0 | 34 | 355.0 | 33 | 227.0 |
|  | 2800 | 6 | 222.0 | 11 | 283.0 | 15 | 406.0 | 11 | 350.0 |
|  | 3800 | 14 | 0 | 9 | 0 | 12 | 0 | 11 | 0 |
| 5 *Lactuca inermis*<br>Forssk.<br>(Asteraceae) | 2300 | 25 | 2.5 | 0 | 00 | 30 | 5.1 | 25 | 3.6 |
|  | 2800 | 12 | 3.5 | 0 | 04 | 9 | 8.7 | 17 | 6.6 |
|  | 3400 | 9 | 0.2 | 0 | 00 | 10 | 3.7 | 9 | 1 |

|  |  |  |  |  |  |  |  |  |  |  |
| --- | --- | --- | --- | --- | --- | --- | --- | --- | --- | --- |
|  |  | 3800 | 10 | 0 | 0 | 00 | 1 | 0 | 11 | 0 |
| 6 | *Senecio burtonii*<br>Hook.f.<br>(Asteraceae) | 2800 | 19 | 0 | 17 | 1.2 | 15 | 14.4 | 19 | 2.9 |
|  |  | 3400 | 2 | 0 | 5 | 0.8 | 1 | 17 | 5 | 5.4 |
| 7 | *Senecio purpureus*<br>L.<br>(Asteraceae) | 2300 | 25 | 0.6 | 20 | 4.3 | 21 | 19.8 | 28 | 11.2 |
|  |  | 2800 | 31 | 1.5 | 25 | 5.6 | 27 | 25.3 | 35 | 18.3 |
|  |  | 3400 | 6 | 0 | 8 | 2.8 | 7 | 22.4 | 6 | 6 |
|  |  | 3800 | 38 | 0.1 | 32 | 1.1 | 22 | 5.2 | 34 | 0.3 |

The hand-pollination experiment followed protocols previously used in Bartoš et al., (2020a,b) based on methods dissected in Dafni et al., (1995). All accessible flower buds on selected plant individuals were bagged with nylon mesh bags before anthesis and monitored until opening. Once flowers opened and stigmas appeared receptive, we applied one of four treatments: autogamy, geitonogamy, outcrossing, and control (Fig. 2A). In the *autogamy* treatment, flowers remained bagged throughout to allow autonomous self-pollination. In the *geitonogamy* treatment, flowers were briefly unbagged and hand-pollinated using pollen from another flower on the same individual, then re-bagged. In the *outcrossing* treatment, pollen from a distant (≥50 m) conspecific individual was applied before re-bagging. *Control* flowers were left unmanipulated and exposed to natural pollination, and they were bagged only after anthesis was complete. After treatment, all flowers remained bagged until collection to prevent seed loss and minimise seed predation.

**Figure 2.**
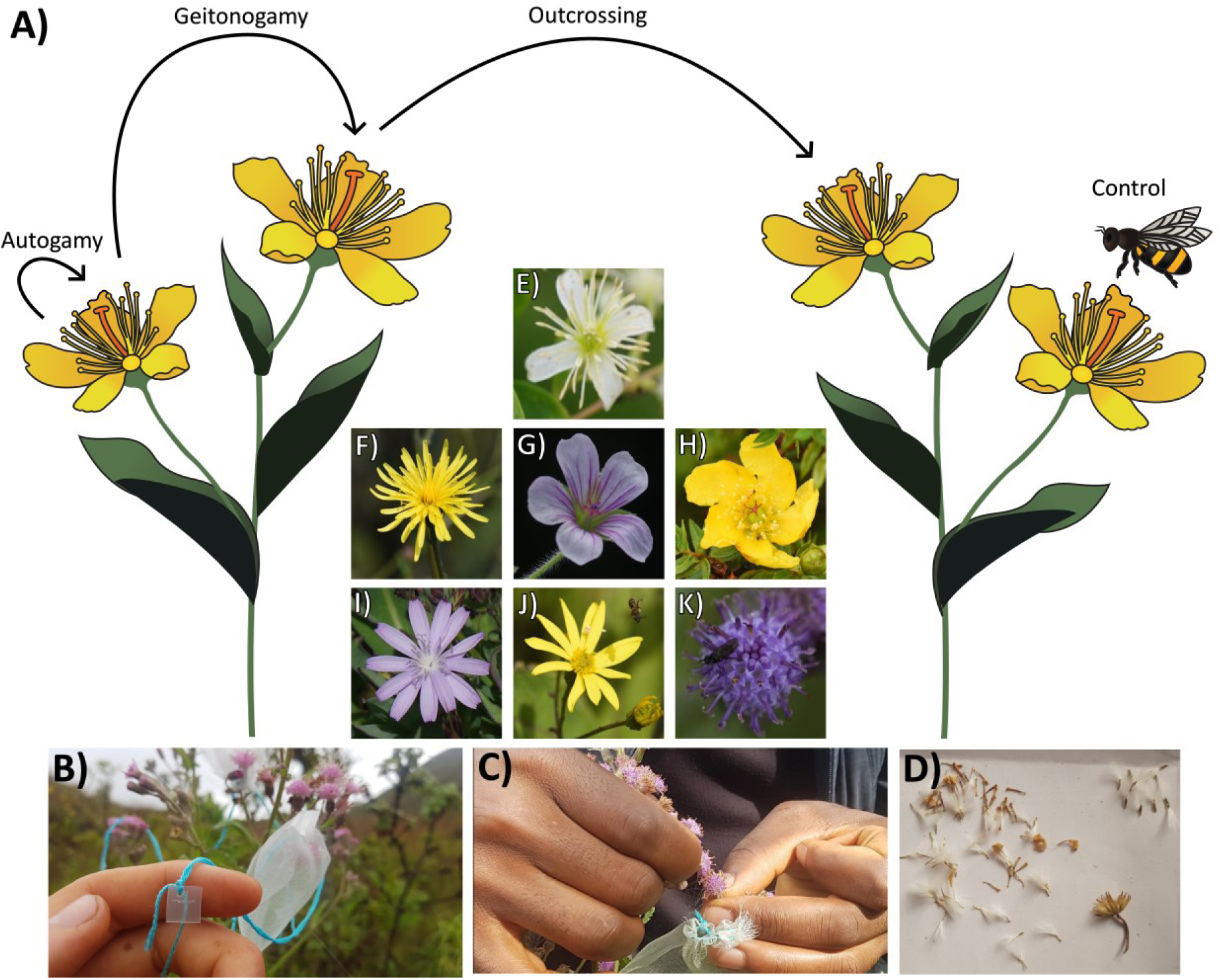
Experimental pollination treatments and studied plant species. A) Schematic representation of the four hand-pollination treatments: autogamy (flowers kept bagged for autonomous selfing), geitonogamy (hand-transferred pollen from other flowers on the same individual), outcrossing (hand-transferred pollen from distant conspecific individuals), and control (flowers exposed to natural pollination). B) Flowers enclosed in mesh bags to exclude visitors and protect treated flowers. C) Hand pollination of experimental flowers. D) Counting seeds from collected inflorescences. E–K) Flowers of the seven focal species: E) *Clematis simensis,* F) *Crepis hypochoeridea,* G) *Geranium arabicum,* H) *Hypericum revolutum,* I) *Lactuca inermis,* J) *Senecio burtonii,* K) *Senecio purpureus*. Credits: Barbora Drozdová (A), Dominik Anýž (B-D), Štěpán Janeček & Petra Janečková (E–K).

For *Geranium arabicum* and *Lactuca inermis*, often only a flower or an inflorescence, respectively, per individual was open at a time, preventing the geitonogamy treatment despite their local abundance. In composite species with sequentially opening florets within a multiple day capitulum (*Crepis hypochoeridea* and *Senecio* spp.), the treatment was repeated daily until all florets had opened; for the purposes of our analyses, each capitulum is referred to as a flower.

All experimental flowers were collected 30–40 days after treatment, dried, and examined under a stereomicroscope to count mature seeds. In composite species, seeds were counted per capitulum. In *Hypericum revolutum*, which produces hundreds of minute seeds in a five-part capsule (Janeček et al., 2007), seeds were spread evenly on a 10 × 10 cm grid and counted within diagonal squares, with total seed number estimated by extrapolation; only intact capsule parts (individual capsule parts were often infested by herbivores or otherwise damaged) were included, and seed set was expressed as the number of seeds per intact capsule part.

### Reproductive indices

Reproductive output under each treatment was quantified from the resulting seed sets. The outcrossing treatment was considered to provide the maximum potential seed set for a flower at a given elevation, because stigmas were saturated with pollen from a distant conspecific individual, thereby minimizing pollen limitation and compatibility constraints. Each species at each elevation was characterised using four reproductive indices: 1/ *natural seed set*, expressed as the mean seed set of control flowers; 2/ *pollen limitation*, calculated as 1 – (control seed set/outcrossing seed set) (Larson and Barrett, 2000); 3/ *autogamy*, calculated as autogamy seed set divided by outcrossing seed set; and 4/ *geitonogamy*, calculated as geitonogamy seed set divided by outcrossing seed set.

When individual-level outcrossing seed set was missing, most often due to loss of treated flowers through herbivory, infestation or abscission before fruit maturation, the species × elevation mean outcrossing seed set was used as a substitute. Pollen limitation values < 0 (i.e. when control seed set exceeded outcrossing seed set) were set to zero. At 3,800 m, *H. revolutum* produced no seeds under any treatment, likely due to unsuitable environmental conditions; at this elevation, the species was therefore retained for analyses of natural seed set but excluded from indices requiring non-zero outcrossing values. In species with multiple flowers/inflorescences per plant, additional buds were often bagged and treated as backups against flower loss; when these flowers matured, their seed sets were also counted, resulting in repeated measurements per individual for some treatments.

### Flower visitors

Flower visitation was observed at all four elevations using continuous 24-hour video recordings of flowering plants with security cameras (VIVOTEK IB8367T with IR night vision), following protocols described in Mertens et al. (2021) and Klomberg et al. (2022). For each focal plant species, seven flowering individuals were recorded per elevation, using plant individuals different from those treated in the hand-pollination experiment but recorded simultaneously with the hand-pollination experiment within the same study sites.

Three visitation metrics were derived from each recorded plant individual. *Visitation frequency* was quantified as the number of visits by any visitor per recorded flower per minute. *Morphospecies richness* was defined as the number of distinct visitor morphospecies, and *functional-group richness* as the number of visitor functional-groups (hoverflies, bees, wasps, birds, beetles, butterflies, moths, and other flies).

### Statistical Analyses

All analyses were performed in R version 4.4.1 (R Core Team, 2024).

To assess how reproductive indices and visitation metrics varied with elevation, we fitted Bayesian generalized linear mixed models (GLMMs) with a consistent hierarchical structure across responses (Korner-Nievergelt et al., 2015; Green et al., 2020). Elevation was included as a categorical fixed effect, whereas species as a random intercept to account for species-specific baselines, and plant individual (nested within species) as an additional random intercept for reproductive indices to account for repeated measurements. Models were fitted using the *brms* package version 2.22.12 (Bürkner, 2017), or, for bounded response variables, using the *ordbetareg* package version 0.8 (Kubinec, 2023). Model distribution (Table 2) was chosen after comparing a set of plausible alternatives identified during the visual inspection (Table S2). Weakly informative priors were used throughout. Model comparison was based on Pareto-smoothed importance sampling leave-one-out cross-validation (PSIS-LOO) and expected log predictive density (ELPD) (Vehtari et al., 2017). When PSIS diagnostics indicated unreliable estimates (Pareto-k > 0.7), we replaced LOO with grouped K-fold cross-validation for the affected response (Vehtari et al., 2017). For each variable, we retained the model with the highest ELPD and checked model adequacy using posterior predictive checks. To evaluate effects of elevation, the selected model was compared with a species-only null model. Elevation was interpreted as influential only when the ELPD difference between the full and null models (ΔELPD) exceeded the difference in its standard error (ΔSE). Following Vehtari et al., (2017, models with |ΔELPD| values smaller than approximately one ΔSE cannot be reliably distinguished in predictive performance, and we therefore favoured the simpler model variant; |ΔELPD| values of about one to two ΔSE were considered weak to moderate evidence; and |ΔELPD| values exceeding two ΔSE represented strong support. Pairwise contrasts among elevations were extracted from posterior draws using the *emmeans* package version 1.11.1 (Lenth, 2017) and regarded as credibly different when their 95% credible intervals excluded zero.

**Table 2.**
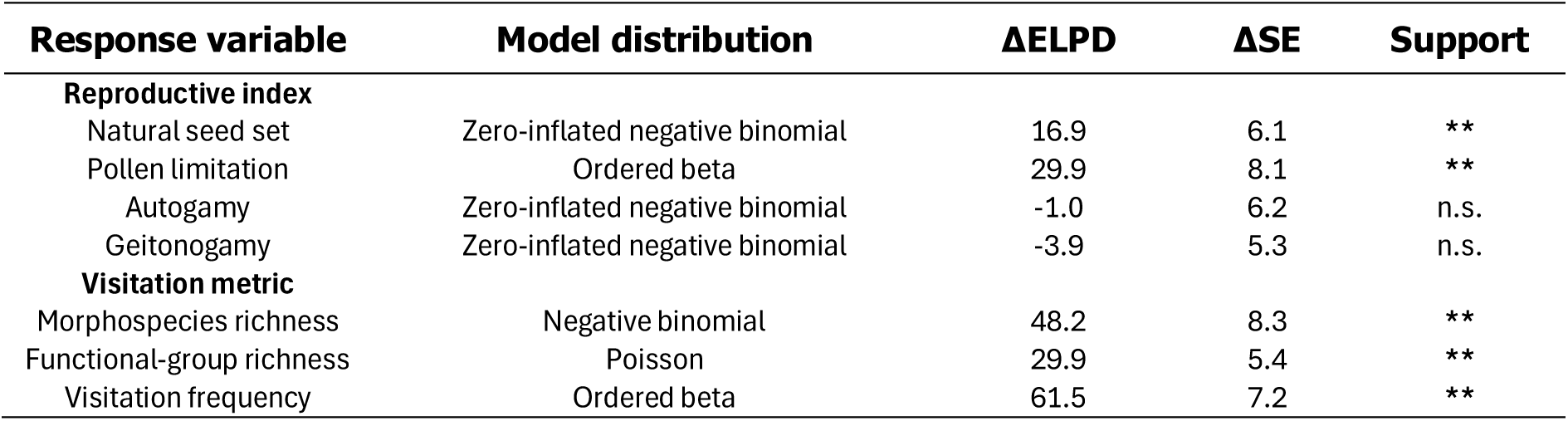
Summary of Bayesian GLMM comparisons testing the effect of elevation on reproductive indices and visitation metrics in Afromontane grasslands on Mount Cameroon. For each response variable, we report the model distribution and the cross-validated difference in expected log predictive density (ΔELPD) between the full model including elevation and the species-only null models, together with the corresponding standard error (ΔSE). Positive ΔELPD values favour the full model, whereas negative ΔELPD values favour the null model. “Support” indicates whether the full model outperformed the null model (|ΔELPD| > 2×ΔSE = strongly supported, “**”; |ΔELPD| ≈ 1–2×ΔSE = weakly to moderately supported, “*”; |ΔELPD| < 1×ΔSE = not supported, “n.s.”).

We then assessed whether variation in reproductive indices could be explained by visitation metrics, using the same hierarchical framework (see Table 3 for model distribution) and the same ΔELPD decision rules. Because reproductive and visitation data originated from different plant individuals, observations were aggregated to species × elevation combinations by pairing each species’ mean reproductive index with its corresponding visitation metric. Morphospecies richness and functional-group richness were strongly correlated (Pearson *r* = 0.80), so to avoid collinearity, only morphospecies richness was retained in these analyses. Visitation predictors were z-transformed, and their across-replicate standard deviations were incorporated using the *brms::me()* function to propagate measurement uncertainty (Bürkner, 2017). Reproductive responses were weighted by a scaled inverse of their within-group standard deviation to account for differing precision in mean seed set estimates. For each reproductive index, we fitted models with all meaningful combinations of visitation predictors as fixed effects, with species included as a random intercept. As in the elevational analyses, predictor effects were interpreted only when the selected model outperformed the species-only null model, and effects were regarded as credibly directional when the 95% credible interval of the regression coefficient excluded zero.

**Table 3.**
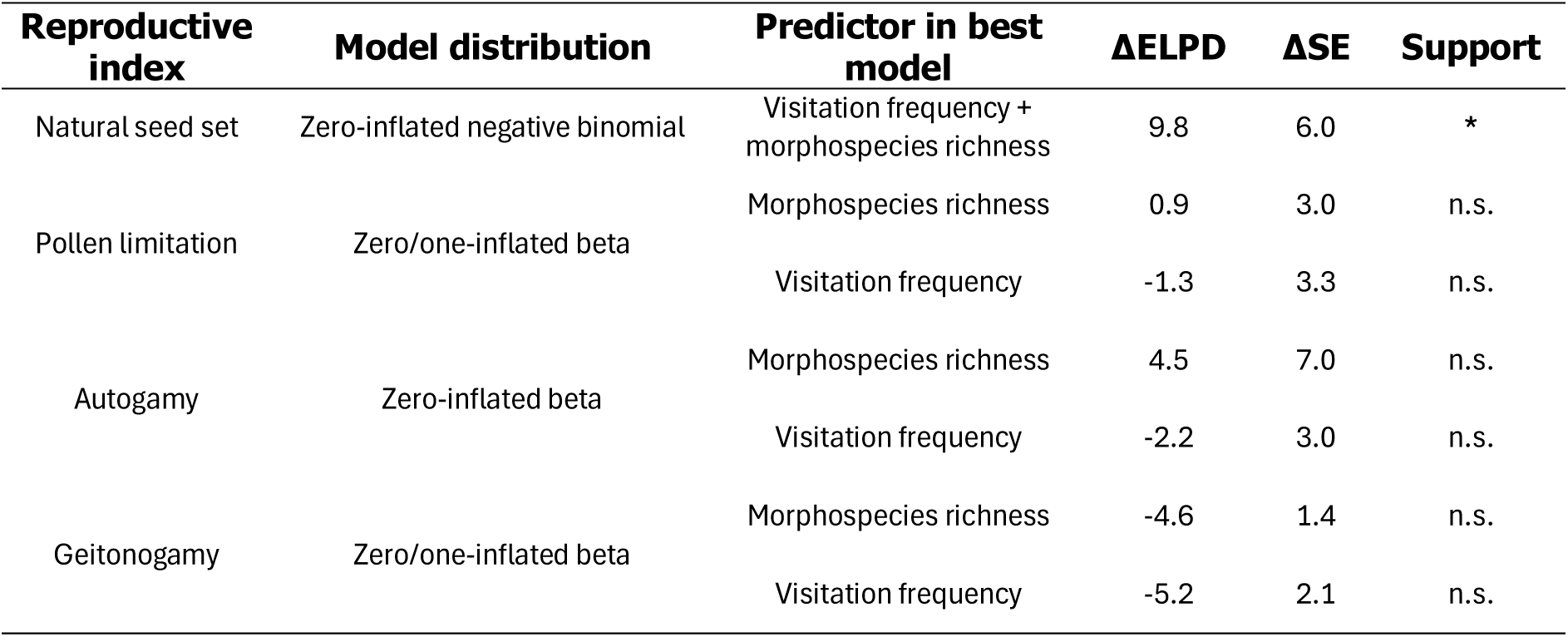
Summary of Bayesian GLMM comparisons testing the effects of visitation metrics on reproductive indices in Afromontane grasslands on Mount Cameroon. For each reproductive index, we report the model distribution and the predictor set in the best cross-validated model (see Table S3 for comparisons across the full candidate set of predictor combinations). ΔELPD is the difference in expected log predictive density between the best models and the species-only null model, together with the corresponding standard error (ΔSE). Positive ΔELPD values favour the predictor (full) model, whereas negative ΔELPD values favour the null (species-only) model. “Support” indicates whether the best model outperformed the null model (|ΔELPD| > 2×ΔSE = predictors strongly supported, “**”; |ΔELPD| ≈ 1–2×ΔSE = weakly to moderately supported, “*”; |ΔELPD| < 1×ΔSE = not supported, “n.s.“).

## Results

We applied hand-pollination treatment to 1,776 flowers at the four studied elevations, of which 1,248 flowers (70%) produced usable seed-set data after losses due to infestation, flower abscission, or technical issues. Plant species were flowering at different numbers of elevations: two occurred at all four elevations, two at three elevations, and three at two elevations (Table 1). *Senecio purpureus* was excluded from visitation analyses at 2,300 m because too few flowering individuals were available for adequate video recording. Floral visitation was quantified from 3,192 hours of video recording, yielding 17,377 recorded visitors, 10,023 of which were identified to 133 morphospecies representing the eight functional groups. Visitation metrics were calculated for each species × elevation combination. Depending on the data completeness for each reproductive index, the resulting sample size for the elevational and visitation-reproduction models ranged from 13 to 20 species × elevation combinations (Table 1).

Bayesian analyses provided strong support for elevational effects on only two reproductive indices, natural seed set and pollen limitation, while no support was found for autogamy and geitonogamy (Table 2, Fig. 3). Natural seed set peaked at 2,800 m and was lower at 2,300 m and 3,400 m, with a pronounced decline at 3,800 m (Table S1, Fig. 3a). Pollen limitation showed the opposite elevational pattern, with substantially higher values at 3,800 m than at any lower site (Table S1, Fig. 3b).

**Figure 3.**
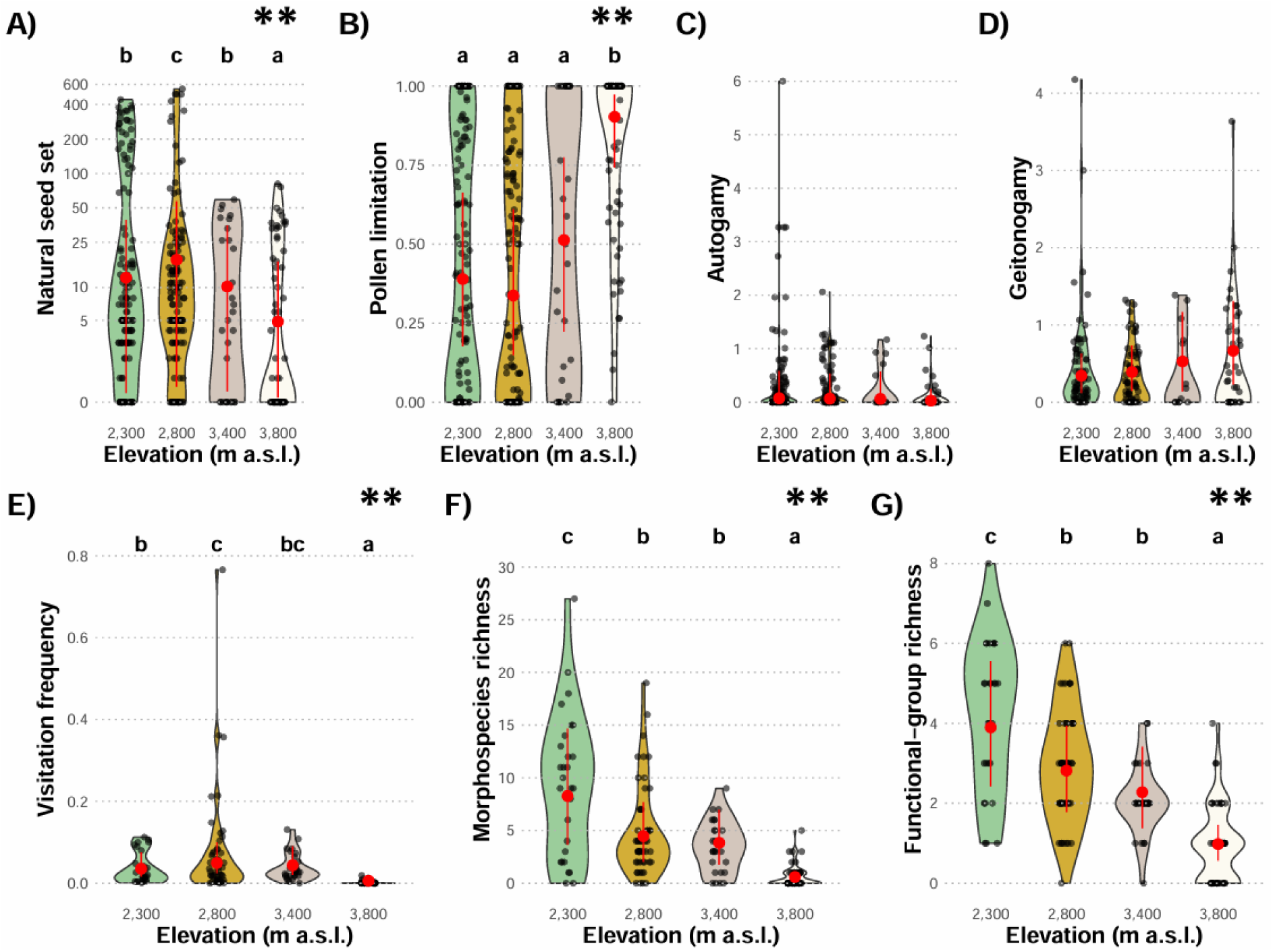
Elevational patterns in reproductive indices (A–D) and visitation metrics (E–G) of plants in Afromontane grasslands on Mount Cameroon. Violin plots show the distribution of observed values, while red dots and vertical lines indicate the marginal means and 95% credible intervals estimated from the corresponding Bayesian GLMM for each response. Natural seed set is shown as the number of seeds per flower or inflorescence, visitation frequency as visits per flower per minute, and visitor richness metrics as numbers of morphospecies and functional groups; the other indices are dimensionless. Letters above violins denote results of posterior pairwise contrasts among elevations, elevations sharing a letter are not credibly different. Asterisks indicate the strength of support for the elevational effect based on cross-validated expected log predictive density (ELPD; see Table 2), with “*” indicating weak to moderate support and “**” indicating strong support for including elevation to the species-only null model.

Individual species exhibited diverse elevational patterns in reproductive indices (Fig. S1). Natural seed set consistently peaked at 2,800 m across all species except *Senecio burtonii*. Pollen limitation showed broadly concordant patterns, with lowest values at 2,800 m and markedly higher values at 3,800 m for the species flowering at this elevation. By contrast, autogamy and geitonogamy showed highly inconsistent elevational patterns (Fig. S1).

All visitation metrics varied with elevation, with broadly similar patterns across indices (Table 2, Fig. 3). Visitation frequency, morphospecies richness, and functional-group richness all declined strongly at 3,800 m, whereas the two mid-elevations showed intermediate and overlapping values (Table S1, Fig. 3).

Among the reproductive indices, only natural seed set showed a supported association with visitation metrics, whereas pollen limitation, autogamy, and geitonogamy did not outperform their species-only null models (Table 3). For natural seed set, the best model included both visitation frequency and morphospecies richness as predictors, outperforming both single predictor models (Table 3; Table S3). The model indicated a clear positive relationship between natural seed set and visitation frequency and a moderate negative relationship between morphospecies richness and natural seed set (Fig. 4).

**Figure 4.**
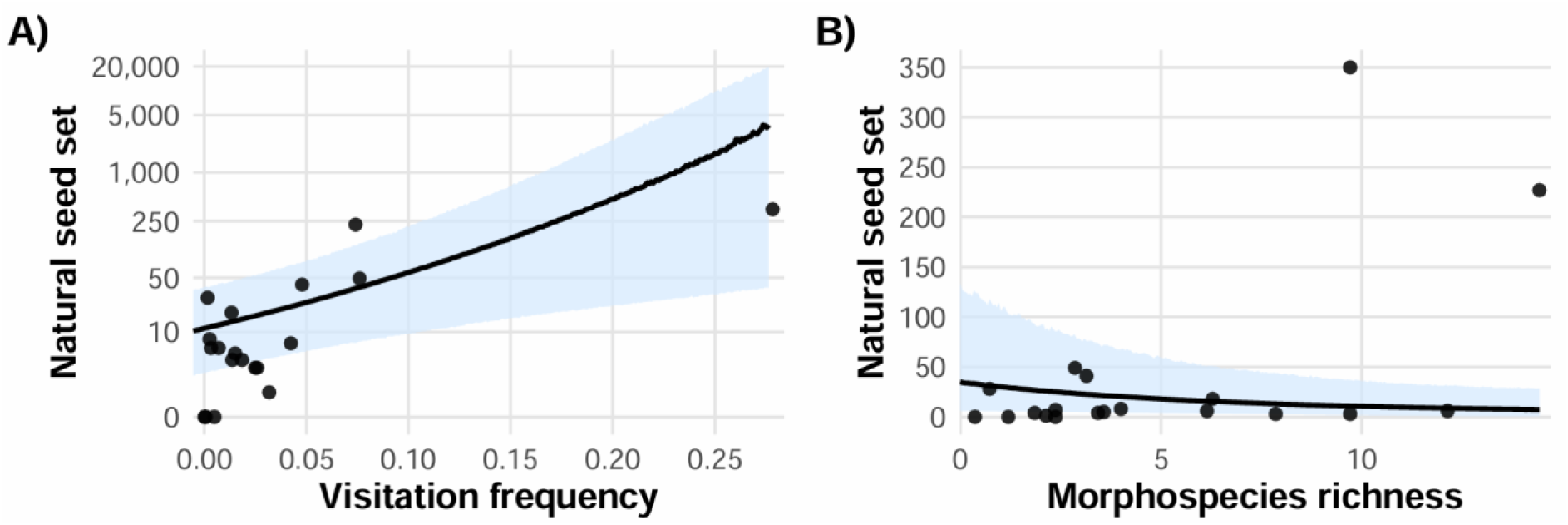
Effects of visitation metrics on natural seed set in Afromontane grasslands on Mount Cameroon, bases on the best supported model by Bayesian GLMM (Table 3). Plots show the marginal effects of A) visitation frequency (visits per flower per minute) and B) morphospecies richness on natural seed set (number of produced seeds per flower/inflorescence). Black lines represent model estimates, with shaded areas showing their 95% credible intervals. See Table 3 for full model summaries, including relationships not supported by the models.

## Discussion

Our study provides a multi-species assessment of intraspecific variation in reproductive success, pollen limitation, selfing, and pollinator visitation along the full elevational gradient of Afromontane grasslands. Across the seven plant species on Mount Cameroon, we found pronounced elevational variation in reproductive success and pollinator visitation, whereas neither autogamy nor geitonogamy showed any consistent elevational change. Natural seed set peaked at lower mid-elevation (2,800 m) and declined steeply towards the summit, where pollen limitation was strongest and visitation frequency and diversity were lowest. All visitation metrics declined towards the highest elevation and explained the elevational patterns in natural seed set, whereas pollen limitation and selfing indices showed no clear relationships with the visitation metrics. Together, these results indicate that reduced pollinator service strongly constrains reproduction at the highest elevation, yet this constraint is not matched by any marked increase in selfing capacity in the focal grassland flora.

### Elevational patterns in reproductive success and pollen limitation

Our results demonstrated that reproductive success and pollen limitation are strongly structured by elevation, with consistently high natural seed set at mid-elevations and a marked decline towards the summit. The overall patterns were consistent across species, fitting our expectations for high elevations where colder, windier and less predictable conditions reduce pollinator diversity, abundance, and activity (Arroyo et al., 1985; Knight et al., 2005; Jiang and Xie, 2020; Mertens et al., 2021; Gaona et al., 2025). The relatively strong pollen limitation detected across the gradient and its pronounced increase towards the summit, is consistent with global evidence that alpine plants often receive insufficient pollen for full seed set (García-Camacho and Totland, 2009; Wagner et al., 2016; Arroyo et al., 2017; Dai et al., 2017; Jiang and Xie, 2020).

The mid-elevation peak in natural seed set and visitation metrics suggests that there is an elevational zone in the montane grasslands where abiotic conditions and pollinator activity jointly favour plant reproduction. In the Afromontane grasslands on Mount Cameroon, the reduced reproductive success at lower elevations may be linked to vegetation structure, as grasslands just above the timberline are locally dominated by a few tall grass species (Janeček and Janečková, 2025) which can reduce both pollinator diversity and the effective floral display of insect-pollinated herbs. The complete failure of *H. revolutum* to produce any seeds at the highest elevation illustrates the severity of abiotic constraints at the upper boundary of the grassland communities. The lack of seed production even when provided with outcross pollen, together with numerous records of two bee morphospecies at this elevation in our study (a functional group previously identified as the primary pollinators of this plant (Bartoš et al., 2015), suggests that unsuccessful reproduction was more likely driven by harsh abiotic conditions than insufficient pollination service at this edge of the gradient. For some plants, such physiological constraints may prevent successful seed production irrespective of pollen availability and thereby amplify the consequences of any shortages in pollination service (Baker, 1972; Guo et al., 2010). Together, these results indicate that elevation-related variation in pollinator service may strengthen or shift the role of abiotic drivers in the elevational patterns of plant reproductive success.

### Elevational patterns in selfing and self-compatibility

In contrast to reproductive success and pollen limitation, intraspecific patterns in selfing showed no detectable responses of both autogamy and geitonogamy to increasingly harsh abiotic conditions and declining pollination service with elevation. This contrasts with our predictions that selfing should become more prevalent where pollinator service is unreliable (Schoen et al., 1996; Busch and Delph, 2012; Barrett et al., 2014). However, studies from other montane systems revealed mixed intraspecific selfing patterns, with some species showing increased selfing at higher elevations (Etcheverry et al., 2008; Seguí et al., 2018; Abdusalam et al., 2020), others showing no consistent elevational trends (Gugerli, 1998; Young et al., 2002; Dai et al., 2017; Black et al., 2019), and some exhibiting even declining selfing along elevation (Wirth et al., 2010).

The relative stability of selfing along the elevational gradient on Mount Cameroon may reflect constraints on the evolution of elevation-specific mating-system adaptations rather than their absence of selective value. Most studied species already exhibited partial self-compatibility at lower elevations, indicating that some degree of reproductive assurance is available without requiring substantial shifts in mating systems at higher sites. This baseline capacity for selfing may buffer populations against declining pollination service and thereby weaken directional selection for further increases in selfing along the gradient. At a broader evolutionary scale, the geologically young age of Mount Cameroon (<3 million years; Wandji et al., 2009), together with frequent volcanic disturbances and increasing anthropogenic pressure (Doležal et al., 2022), may have limited the time and stability required for consistent elevation-related selection on mating systems. In addition, the limited extent and isolation of Afromontane grasslands likely constrain effective size and genetic diversity of populations, increasing the role of genetic drift and reducing the efficiency of selection, while the close spatial proximity of grassland sites along the gradient may promote gene flow that counteracts divergent selection and prevents the fixation of local adaptations. Together, “baseline” self-compatibility, limited evolutionary time, frequent disturbance, small population sizes, and ongoing connectivity among populations may collectively constrain the emergence of consistent elevational differentiation in selfing and self-compatibility, even in the presence of strong elevational gradients in pollinator service.

### Pollinator visitation and elevational variation in seed set

The positive relationship between natural seed set and pollinator visitation in our study suggests that elevational differences in plant reproductive success are associated with variation pollination service, reflected jointly in visitor activity and diversity (because functional-group richness was closely correlated with morphospecies richness, we interpret them together). Similar patterns have been reported from other montane systems, where harsher conditions at higher elevations reduced pollinator activity and/or diversity, with consequences for plant reproduction (Arroyo et al., 1985; Totland, 1997; Ramos-Jiliberto et al., 2010; Wu et al., 2019). In contrast, the absence of clear relationships between visitation metrics and pollen limitation indicates that pollination service alone cannot fully explain elevational patterns in reproductive success, highlighting the role of physiological constraints under harsh high-elevation conditions. At the same time, this lack of association may partly reflect our data limitations, as pollen limitation integrates multiple processes beyond visitation frequency, including pollen quality, heterospecific pollen deposition, post-pollination resource or developmental constraints, all of which can decouple it from simple visitation metrics (Knight et al., 2005). Finally, the absence of relationships between visitation metrics and selfing indices is consistent with the lack of elevational patterns in selfing in our study. Autogamy and geitonogamy largely reflect other plant adaptations, such as floral morphology, and evolutionary history of inbreeding depression, and therefore do not necessarily respond to short-term variation in pollinator activity or diversity (Willmer, 2011; Busch and Delph, 2012; Barrett et al., 2014).

### Implications

Our findings place Afromontane grasslands within broader patterns reported from other, mostly temperate and subtropical, mountain systems, while also highlighting some divergences. The strong decline in seed set and pollinator visitation towards the summit corroborates broader evidence that tropical montane plants are strongly constrained by the combined effects of abiotic stress and reduced pollination service near the upper vegetation limits (Arroyo et al., 1985; Jiang and Xie, 2020). In our system, summit populations experience pronounced pollen limitation and very low seed production despite measurable partial self-compatibility in several species, indicating that selfing provides only limited buffering against declining pollination. This suggests that the highest-elevation populations may function as reproductive sinks, relying on seed input from lower sites rather than sustained local populations adapted to the harsh conditions.

Tropical Afromontane ecosystems are currently undergoing rapid climatic and land-use changes (Mata-Guel et al., 2023; Abera et al., 2024), and high-elevation communities are expected to be among the particularly sensitive to shifts in pollinator availability and phenology (Hegland et al., 2009; Trunschke et al., 2024). Within this context, our results imply that upslope range shifts under warming will not necessarily result in successful reproduction if plants track climate into zones where pollinator service, other physiological limits, or their interaction remain unfavourable. Importantly, elevational responses of plants and their pollinators may be asynchronous, leading to spatial or temporal mismatches that further constrain reproduction even where climate becomes suitable for plant persistence. Together, these processes highlight the vulnerability of isolated high-elevation plant communities to ongoing environmental change and human disturbance, while also underscoring the need to protect pollinator assemblages to maintain reproductive functioning under future climates.

## Acknowledgements

We are grateful to Kum Peter Abigha, Francis Teke Mani, Joseph Ekema, Paul Orock Agbor, Frederick Bowen Raboya, Lucas Lyonga Molua, and Marcus Mokake Njie for their assistance during the field work; to Charlotte Bowin and Mercy Murkwe for their help with processing of video recordings; to Eric B. Fokam for his help with permits and other logistics; to Barbora Drozdová for illustrating the experiment design in Fig. 2; and to the MCNP staff for all their assistance. We used ChatGPT (model 5.1; OpenAI) for English proofreading. This study was conducted under authorisations issued by the Ministries of Forestry and Wildlife and of Research and Innovations of the Republic of Cameroon. The research was funded by the Czech Science Foundation (21-24186M). OM was funded the Charles University Research Centre program (UNCE/24/SCI/006), and by the Institutional Support for Science and Research of the Ministry of Education, Youth and Sports of the Czech Republic. Computational resources were provided by the e-INFRA CZ project (ID:90254), supported by the Ministry of Education, Youth and Sports of the Czech Republic.

