## Supplementary material for "Elevational patterns in plant mating systems and pollen limitation in Afrotropical montane grasslands"

***List of supplementary materials:***

Table S1. Pairwise posterior contrasts among elevations for reproductive indices and visitation metrics analysed in the Bayesian GLMMs for Afromontane grasslands on Mount Cameroon.

Table S2. Summary of candidate model distributions evaluated for each response variable in the Bayesian GLMMs for Afromontane grasslands on Mount Cameroon.

Table S3. Summary of candidate Bayesian GLMMs testing the effects of visitation metrics on reproductive indices in Afromontane grasslands on Mount Cameroon.

Figure S1. Intraspecific variation in natural seed set along the elevational gradient on Mount Cameroon.

Figure S2. Intraspecific variation in pollen limitation along the elevational gradient on Mount Cameroon.

Figure S3. Intraspecific variation of the autogamy index along the elevational gradient on Mount Cameroon.

Figure S4. Intraspecific variation of the geitonogamy index along the elevational gradient on Mount Cameroon.

**Table S1. Pairwise posterior contrasts among elevations for reproductive indices and visitation metrics analysed in the Bayesian GLMMs for Afromontane grasslands on Mount Cameroon.** For each response variable, the table reports estimated differences between elevation pairs (Est.) with their 95% credible intervals (CI_95), derived from posterior draws using emmeans. Contrasts were interpreted as credibly different when the 95% credible interval excluded zero, corresponding to the letter-grouping summaries displayed above violin plots in Fig. 3.

| Compared elevations | Natural  seed set | | Pollen limitation | | Geitonogamy | | Morphospecies richness | | Functional-group richness | | Visitation frequency | |
| --- | --- | --- | --- | --- | --- | --- | --- | --- | --- | --- | --- | --- |
|  | **Est.** | **CI_95** | **Est.** | **CI_95** | **Est.** | **CI_95** | **Est.** | **CI_95** | **Est.** | **CI_95** | **Est.** | **CI_95** |
| 2,300 – 2,800 | -0.37 | [-0.57, -0.13] | 0.22 | [-0.29, 0.70] | -0.14 | [-0.58, 0.25] | 0.62 | [0.28, 0.96] | 0.32 | [0.05, 0.57] | -0.35 | [-0.7, -0.01] |
| 2,300 – 3,400 | 0.18 | [-0.29, 0.67] | -0.51 | [-1.34, 0.31] | -0.45 | [-1.22, 0.29] | 0.77 | [0.29, 1.23] | 0.53 | [0.15, 0.92] | -0.22 | [-0.68, 0.22] |
| 2,300 – 3,800 | 0.91 | [0.36, 1.47] | -2.69 | [-3.57, -1.83] | -0.66 | [-1.31, -0.07] | 2.63 | [2.13, 3.14] | 1.38 | [0.99, 1.74] | 1.87 | [1.42, 2.33] |
| 2,800 – 3,400 | 0.55 | [0.09, 0.99] | -0.73 | [-1.51, 0.10] | -0.30 | [-1.02, 0.41] | 0.15 | [-0.23, 0.51] | 0.21 | [-0.12, 0.53] | 0.14 | [-0.2, 0.5] |
| 2,800 – 3,800 | 1.28 | [0.72, 1.80] | -2.92 | [-3.82, -2.10] | -0.51 | [-1.12, 0.09] | 2.01 | [1.57, 2.45] | 1.06 | [0.73, 1.4] | 2.22 | [1.79, 2.643 |
| 3,400 – 3,800 | 0.74 | [0.26, 1.22] | -2.18 | [-3.13, -1.35] | -0.22 | [-0.95, 0.52] | 1.86 | [1.36, 2.35] | 0.85 | [0.46, 1.24] | 2.09 | [1.61, 2.55] |

**Table S2. Summary of candidate model distributions evaluated for each response variable in the Bayesian GLMMs for Afromontane grasslands on Mount Cameroon.** For each response variable, all plausible distributions preselected during visual inspection are reported. Columns show the expected log predictive density (ELPD) and the difference from the best-performing distribution (ΔELPD) with its standard error (ΔSE). “Support” indicates the relative performance of each distribution compared with the best one (|ΔELPD| > 2×ΔSE = strongly different, “**”; |ΔELPD| ≈ 1–2×ΔSE = weakly to moderately different, “*”; |ΔELPD| < 1×ΔSE = indistinguishable, “n.s.”).

| Response variable | Model distribution | ELPD | ΔELPD | ΔSE | Support |
| --- | --- | --- | --- | --- | --- |
| Natural seed set | **Zero-inflated negative binomial** | **-1191.97** | **0** | **0** | **best model** |
|  | Negative binomial | -1251.35 | -59.4 | 13.9 | ** |
|  | Poisson | -2531.68 | -1339.7 | 374.6 | ** |
|  | Zero-inflated Poisson | -2662.33 | -1470.4 | 404.1 | ** |
| Autogamy | **Zero-inflated negative binomial** | **-795.123** | **0** | **0** | **best model** |
|  | Negative binomial | -811.423 | -16.3 | 10.4 | * |
|  | Zero-inflated Poisson | -1087.11 | -291.9 | 120.6 | ** |
|  | Poisson | -1372.45 | -577.3 | 154.8 | ** |
| Geitonogamy | **Zero-inflated negative binomial** | **-866.399** | **0** | **0** | **best model** |
|  | Negative binomial | -931.03 | -64.6 | 8.1 | ** |
|  | Zero-inflated Poisson | -983.099 | -116.7 | 41.7 | ** |
|  | Poisson | -2606.13 | -1739.7 | 506.2 | ** |
| Morphospecies richness | **Negative binomial** | **-321.911** | **0** | **0** | **best model** |
|  | Zero-inflated negative binomial | -322.337 | -0.4 | 1.5 | n.s. |
|  | Zero-inflated Poisson | -339.091 | -17.2 | 8.7 | ** |
|  | Poisson | -345.658 | -23.7 | 9.5 | ** |
| Functional-group richness | **Poisson** | **-235.89** | **0** | **0** | **best model** |
|  | Zero-inflated Poisson | -236.907 | -1.0 | 0.2 | ** |
|  | Zero-inflated negative binomial | -238.267 | -2.4 | 0.3 | ** |

**Table S3. Summary of candidate Bayesian GLMMs testing the effects of visitation metrics on reproductive indices in Afromontane grasslands on Mount Cameroon.** For each reproductive index, all fitted combinations of visitation predictors (visitation frequency, morphospecies richness) are reported. Columns show the expected log predictive density (ELPD) from K-fold cross-validation and the difference from the best-performing model (ΔELPD) with its standard error (ΔSE). “Support” indicates the relative performance of each model compared with the best model (|ΔELPD| > 2×ΔSE = strongly different, “**”; |ΔELPD| ≈ 1–2×ΔSE = weakly to moderately different, “*”; |ΔELPD| < 1×ΔSE = indistinguishable, “n.s.”).

| Reproductive index | Predictor combination | ELPD | ΔELPD | ΔSE | Support |
| --- | --- | --- | --- | --- | --- |
| Natural seed set | visitation frequency + morphospecies richness | -110.824 | 0 | 0 | **best model** |
|  | morphospecies richness | -118.838 | -8.0 | 3.8 | ** |
|  | visitation frequency | -119.499 | -8.7 | 2.9 | ** |
| Pollen limitation | morphospecies richness | -7.31184 | 0 | 0 | **best model** |
|  | visitation frequency | -9.54497 | -2.2 | 2.9 | **n.s.** |
|  | visitation frequency + morphospecies richness | -10.2669 | -3.0 | 2.7 | * |
| Autogamy | visitation frequency | -13.4787 | 0 | 0 | **best model** |
|  | visitation frequency + morphospecies richness | -18.6854 | -5.2 | 5.4 | **n.s.** |
|  | morphospecies richness | -20.2233 | -6.7 | 6.7 | **n.s.** |
| Geitonogamy | morphospecies richness | -2.001 | 0 | 0 | **best model** |
|  | visitation frequency | -2.5932 | -0.6 | 2.8 | **n.s.** |
|  | visitation frequency + morphospecies richness | -4.7319 | -2.7 | 1.3 | **n.s.** |

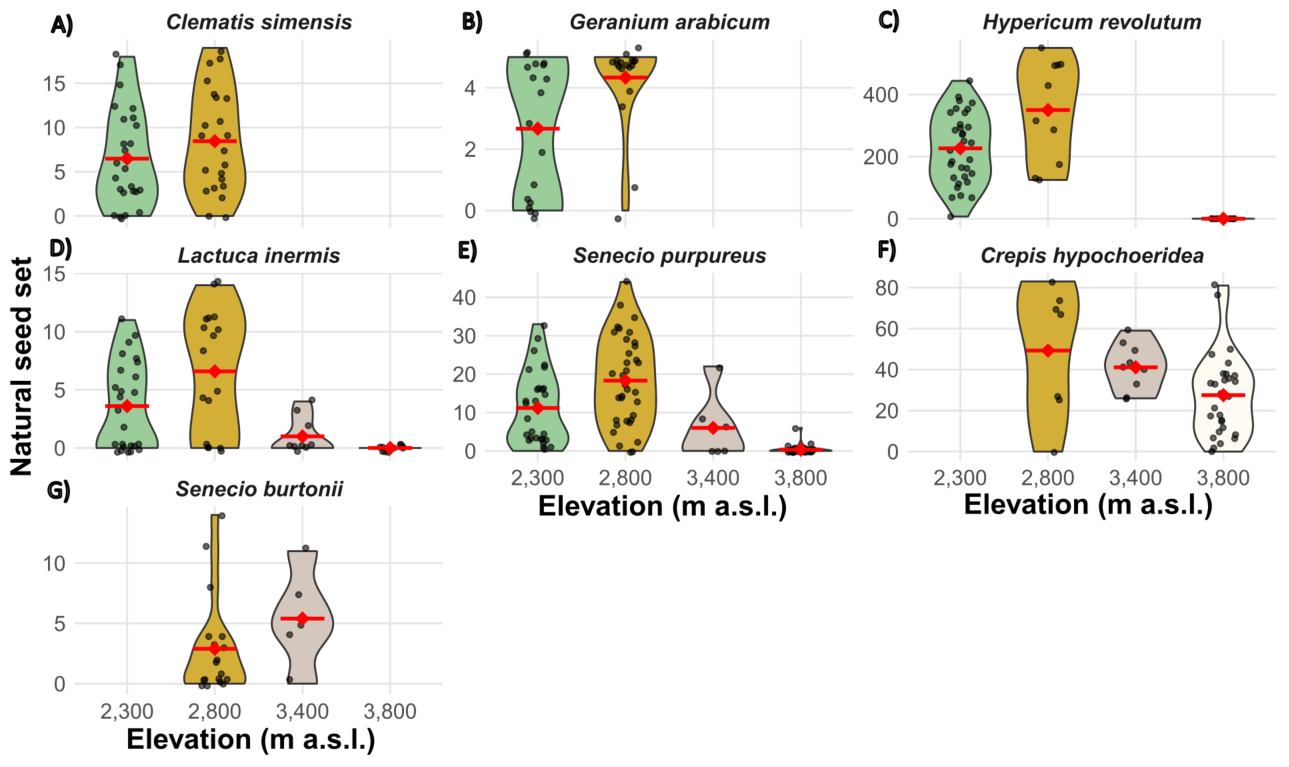

**Figure S1. Intraspecific variation in natural seed set along the elevational gradient on Mount Cameroon.** Violin plots show, for each species, the distribution of observed values, while red dots with horizontal bars indicate mean values per elevation.

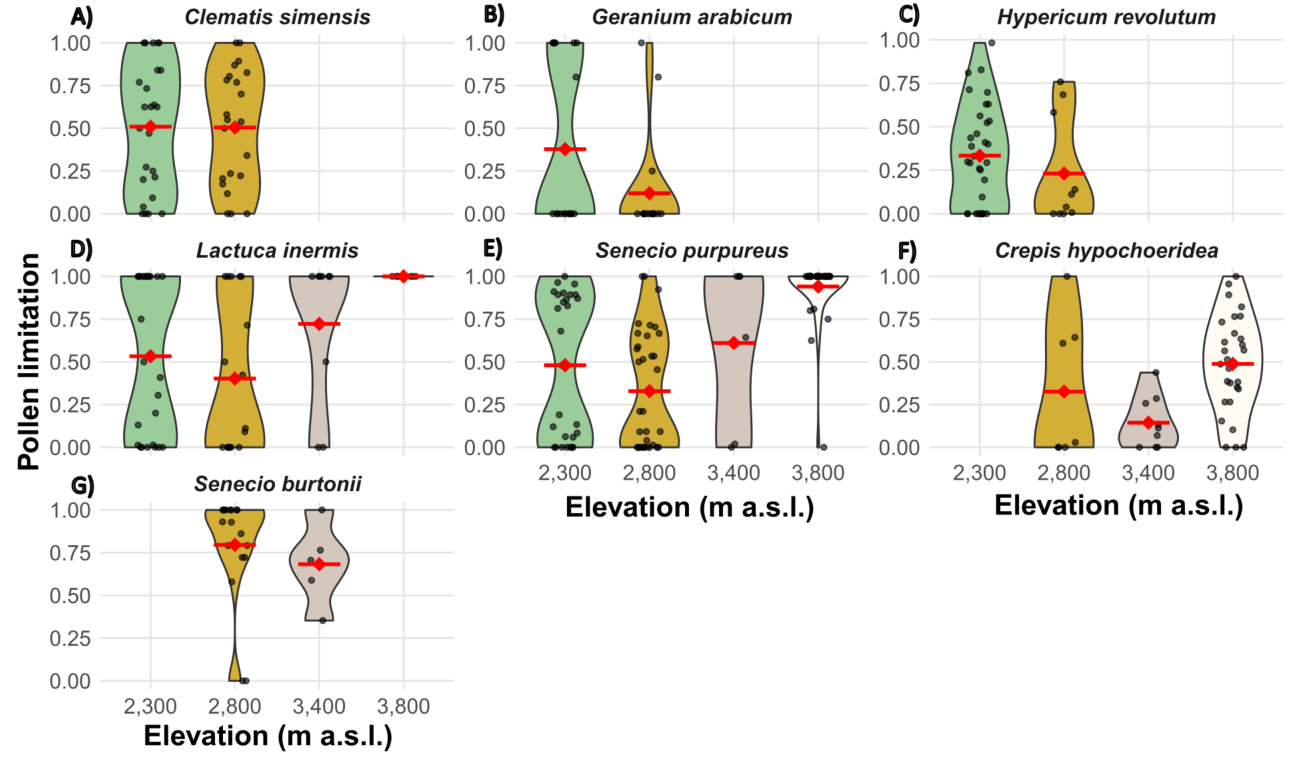

**Figure S2. Intraspecific variation in pollen limitation along the elevational gradient on Mount Cameroon.** Violin plots show, for each species, the distribution of observed values, while red dots with horizontal bars indicate mean values per elevation.

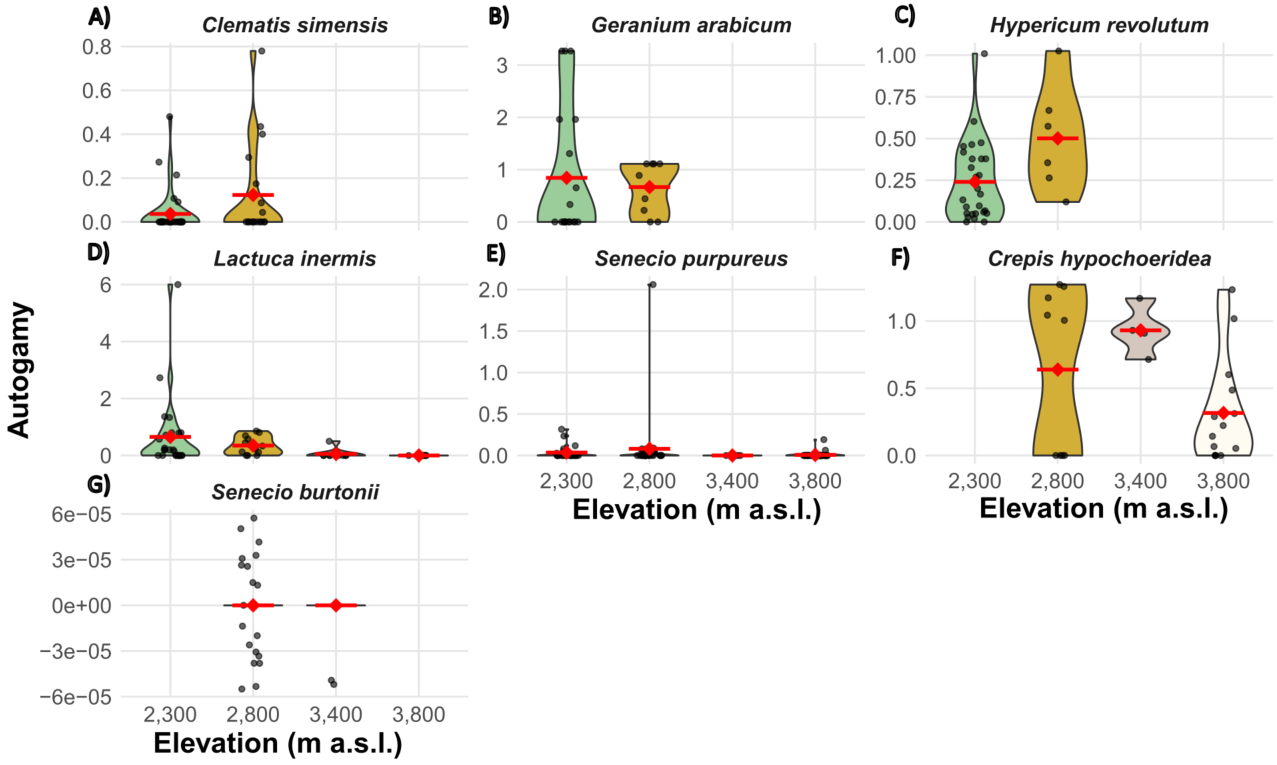

**Figure S3. Intraspecific variation of the autogamy index along the elevational gradient on Mount Cameroon.** Violin plots show, for each species, the distribution of observed values, while red dots with horizontal bars indicate mean values per elevation.

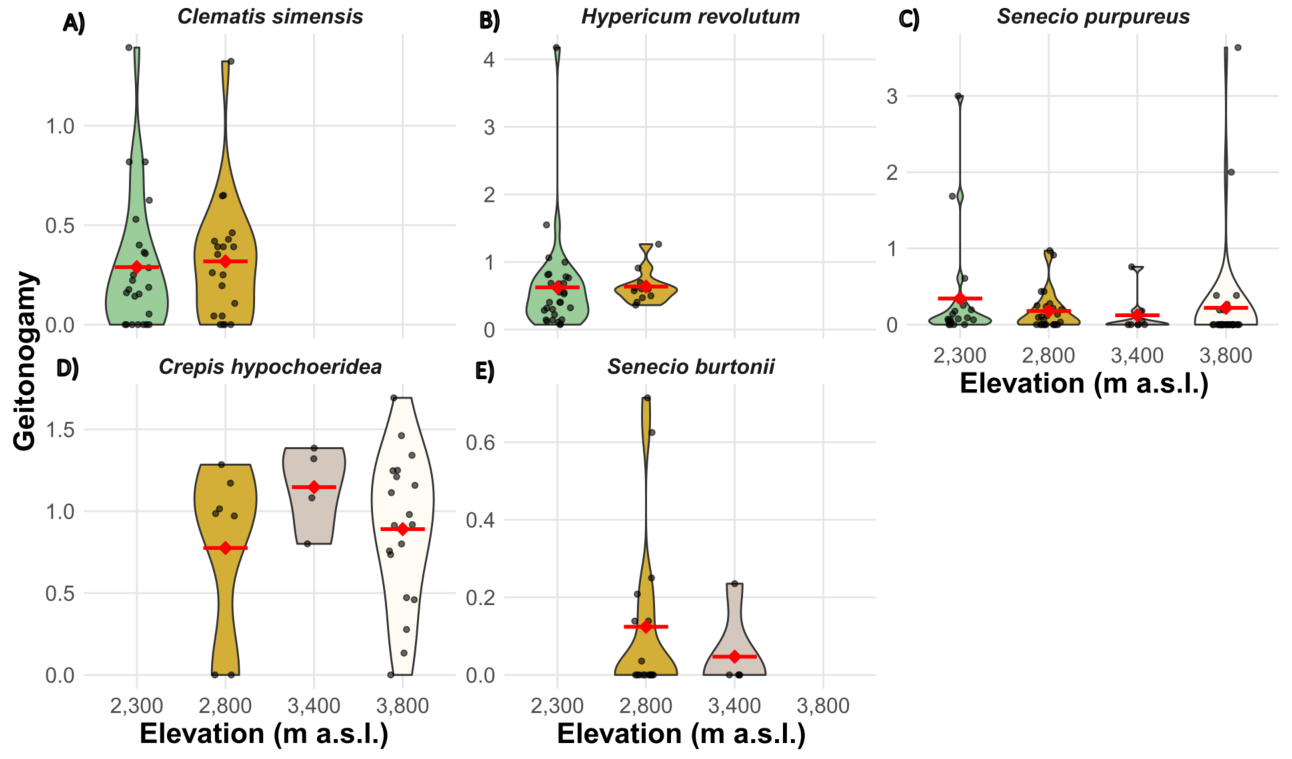

**Figure S4. Intraspecific variation of the geitonogamy index along the elevational gradient on Mount Cameroon.** Violin plots show, for each species, the distribution of observed values, while red dots with horizontal bars indicate mean values per elevation.
